# Prior visual context accelerates the emergence of object representations

**DOI:** 10.64898/2026.10.15.759465

**Authors:** Alex Clarke, Ye Gu, Alexandra Krugliak

## Abstract

Our knowledge about how the world is structured allows for predictions about what objects we might see next. Visual context is known to impact the perception of subsequent objects, but we don’t know what context does to the mental representation of those objects. Using MEG, we tested how a scene context changes the neural representation of visual objects, and whether the context simply enhances expected object representations, or if it acts to reorganise the temporal trajectory of object information in the brain. Using temporal generalisation decoding, we show that multiple dimensions of semantic information are shifted up to 200 ms earlier when the object and the prior context are congruent, and this temporally-shifted decoding was related to behavioural object recognition speed. This suggests that prior context can warp the temporal trajectory through which object representations are accessed and accelerate the emergence of semantic object representations in the mind.

## Introduction

Inferring the meaning of our sensory experiences is arguably at the core of cognition. Semantic object recognition – understanding the meaning of visual objects, feels effortless and almost instantaneous. Studies using time-resolved methods have charted the time-course of object information, providing broad evidence for the initial coding of low-level visual features, followed by superordinate category and higher-level visual features, before semantic object properties (Bankson et al., 2018; Carlson et al., 2013; Cichy et al., 2014; Clarke et al., 2013, 2015, 2018; Jozwik et al., 2016, 2023; Kaiser et al., 2016; Leonardelli & Fairhall, 2022; von Seth et al., 2023). However, these approaches fail to take into consideration how the prior knowledge we have about the context might shape the unfolding dynamics of object perception. In the real world, objects don’t exist in isolation, but are part of an extended spatiotemporal experience. Traditional accounts of object recognition say little about how the preceding context might modulate the dynamics underlying object recognition, but this is vital if we are to understand real-world experiences. Here we assess if, and how, the temporal dynamics of object information are influenced by the relationship between the object and the context within which the object is experienced. This is tested using MEG recordings for objects preceded by congruent or incongruent visual scenes alongside a temporal generalisation approach to neural decoding (King & Dehaene, 2014) to reveal how object information is flexibly shifted in time.

Our knowledge about how the world is structured allows for predictions about what objects we might see next. It has been widely shown that object recognition speed and accuracy is facilitated when objects appear within, or following, a congruent context (Bar, 2004; Biederman et al., 1982; Davenport & Potter, 2004; Gu et al., 2025; Joubert et al., 2007; Oliva & Torralba, 2007; Palmer, 1975). A large body of research using time-sensitive measures, such as EEG and MEG, shows that neural responses during the processing of objects are influenced by the context they’re experienced within, where activity beyond 200 ms is modulated by the relationship between an object and a visual scene, with a larger response magnitude for incongruent trials (Coco et al., 2020; Ganis & Kutas, 2003; Gu et al., 2025; Kumar et al., 2021; Lauer et al., 2018; Mudrik et al., 2014; Võ & Wolfe, 2013). Such modulations are typically seen in a fixed time-window, and have been linked to accessing semantic information about the object, integrating object and contextual information, and detecting semantic violations (Eddine et al., 2024; Henderson & Hollingworth, 1999; Kutas & Federmeier, 2011; Võ, 2021). However, beyond a modulation of response amplitude, a more fundamental question is how context changes the temporal flow of object information.

One mechanism that may help frame the problem is predictive coding, where the brain predicts upcoming sensory signals based on information gleaned from the prior context (de Lange et al., 2018; Eddine et al., 2024; Peelen et al., 2024; Press et al., 2020; Trapp & Bar, 2015). One account, the opposing process theory (Press et al., 2020), argues that visual perception is initially more sensitive to expected information, and when the stimulus does not conform to such expectations, a later boost of the incongruent information occurs.

Overall, this suggests that representations of object information that aligns with those predictions will be altered, but may manifest through either enhanced neural representations, or a temporally shifted representation that is brought forward in time. While the former appears to converge with a temporally-fixed modulation, the latter suggests that the temporal evolution of object information is more flexible, whose trajectory through representational states is conditioned on the prior context.

Here, we address this issue by determining whether different types of object information are changed depending on the relationship to the prior scene – two semantic properties, real-world size and an objects contextual category; and one image-based perceptual dimension, object shape. MEG data were collected during an object recognition paradigm where each object was preceded by a congruent scene, an incongruent scene, or a scrambled image (no context condition; Figure 1). Critically, the same objects were viewed in all context conditions, and the scene was temporally separated from the object, thus allowing us to study changes in object processing due to the mnemonic impact of the scene, rather than the ongoing perception of the scene.

**Figure 1.**
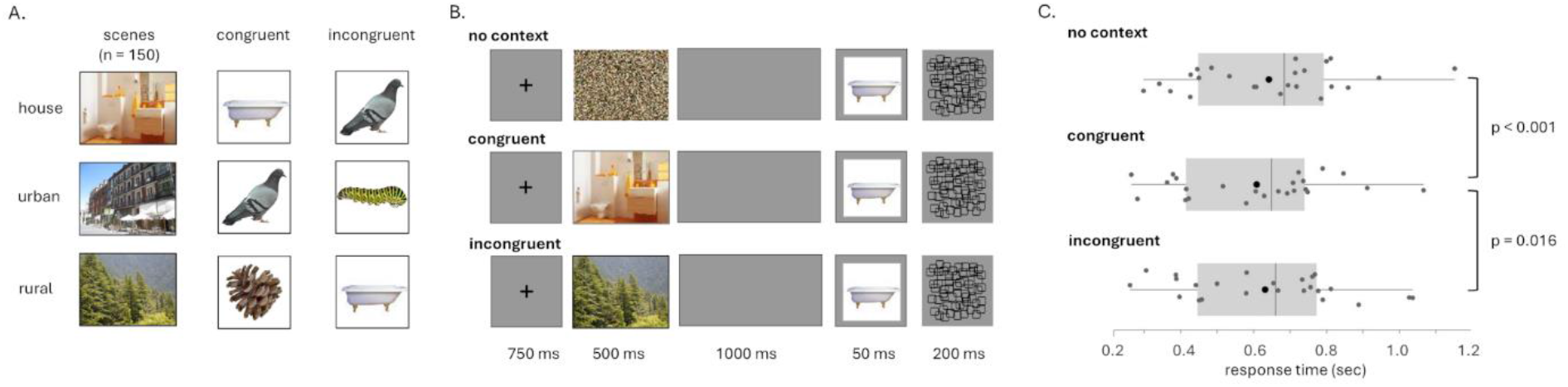
Paradigm and behavioural results. a) Example images of each scene type alongside the congruent and incongruent objects they are paired with. b) Example trail from each condition, where an object is preceded by a scrambled image (no context), a visual scene that is congruent with the object, or a visual scene that is incongruent with the object. Each object appears in each condition, and participants pressed a button when they knew the identity of the object. c) Behavioural responses showing faster response speeds to objects when they followed a congruent scene compared to incongruent or no-context (scrambled) trials.

We look to adjudicate between three plausible hypotheses for how prior context might change object information. First, we might see no impact of context on the coding of object information, as predicted by accounts where objects and scenes are coded independently and integrated at a late, post-recognition stage (Henderson & Hollingworth, 1999). Second, it might be that a congruent object-scene relationship leads to enhanced object decoding without a change in the temporal dynamics, similar to modulation patterns seen in EEG components. Finally, a third plausible hypothesis is that a congruent visual scene prior to the object results in a temporal shift in object information, such that object information coded when no context is present is brought forward in time by the mnemonic influence of the congruent scene. We test between these hypotheses using temporal generalisation decoding of object information. Our critical analyses trained object classifiers on the MEG signals from the no context condition, and used objects from the congruent or incongruent conditions as test trials. Temporal generalisation decoding then allows us to assess if object information is shifted in time for the congruent and incongruent trials relative to the no context condition, and if the impact is specific for semantic, in contrast to image-based properties.

## Results

MEG signals were collected during an object recognition paradigm where participants pressed a button when they recognised the identity of the briefly presented object (Figure 1b). Objects were presented in isolation from the background, but were preceded by a visual context that could be a more congruent scene, an incongruent scene or a scrambled image to create a no-context condition. Each of the 150 objects were presented three times, once in each of the conditions, meaning each of the 150 different scene images appeared with a congruent and an incongruent object. The scene and object were temporally separated so that the visual scene could provide a pre-existing context for the object, more closely mirroring real-life experiences than paradigms where scenes and objects appear simultaneously. Further this means that object decoding performance will not be directly influenced by the joint perception of the scene.

### Object-scene congruency facilitates recognition speed

To assess if the scene-object relationship influenced behavioural recognition speeds, a repeated measures ANOVA with factor scene type (congruent, incongruent, no context) revealed a significant difference in response times (F = 8.46, df = 2,46, p < 0.001). Further analyses of response times showed that participants were faster to recognise objects when they followed a congruent scene (RT = 608 ms, SD = 208 ms) compared to an incongruent scene (RT = 632 ms, SD = 217 ms ; t(23)=2.77, p = 0.016) or compared to the no-scene condition (RT = 642 ms, SD = 211 ms ; t(23)=4.02, p < 0.001; Figure 1c). No differences in response times were seen between incongruent trials and no-scene trials (t(23)=1.25, p = 0.22). In agreement with past research, this suggests that the prior congruent scene facilitated access to object information, and might indicate that recognition following an incongruent scene is more closely aligned with object recognition when no context is present.

### Object information without context

Before asking whether the dynamics of object information is changed due to the prior scene context, we sought to evaluate if our three types of information about objects could be decoded when no context is present – image-shape, real-world size, and contextual category information. Using the MEG signals from the object presentation period, we used a temporal generalisation approach to decoding, where classifiers were trained to distinguish between objects that belong to different categories; image-shape classifiers were trained to distinguish between images where the object was horizontally orientated and images where the object was vertically orientated, real-world size classifiers were trained to distinguish between big and small objects, and contextual category classifiers between household objects, street objects and rural objects (Figure 2a). In all cases, 10-fold cross-validation procedures were used, with decoding accuracy averaged across all 10-folds.

**Figure 2.**
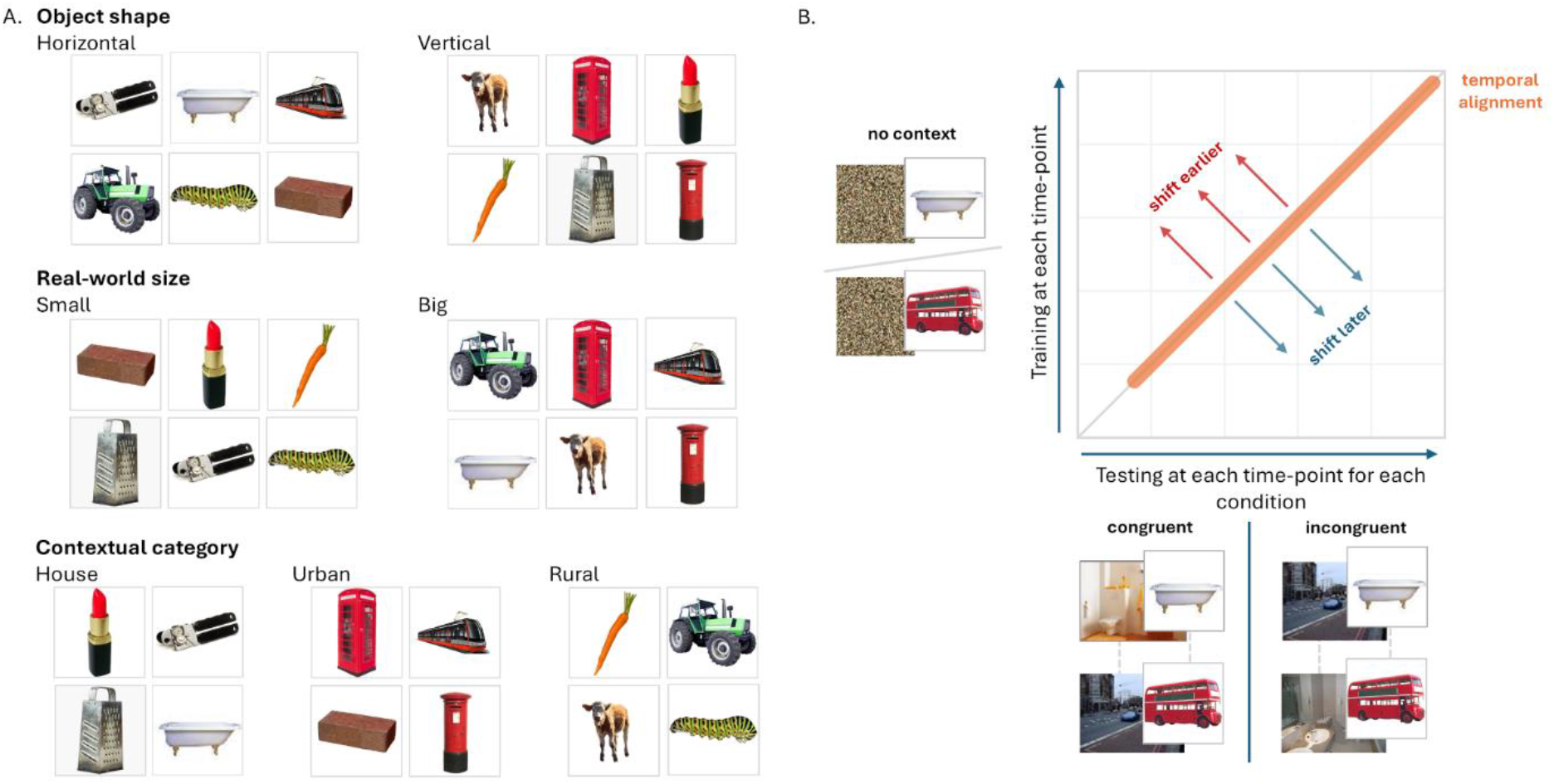
Decoding of object information. a) Illustration of how objects were grouped to decode different object properties from the same set of objects. Twelve example objects are shown grouped in different ways for object shape, real-world size and contextual category decoders. b) MEG decoding of object information across conditions. MEG data from the no-context trials are used to train a classifier. This classifier is then used to decode object information from objects following a congruent/incongruent scene. Decoding along the diagonal means the across-condition classifier decodes object information for the same time-point used for training (temporally aligned). If object information is seen earlier in time compared to the no-context condition then decoding will be present in the upper triangle, and if object information is seen later in time compared to the no-context condition then decoding will be present in the lower triangle. Example images illustrate contextual category decoding.

Classifiers were trained at one time point and tested at every time point across the object epoch. This allowed us to determine whether representations are stably coded over time, or more dynamically changing (King & Dehaene, 2014). For instance, object information could be coded at several different time-points, one after the other, but for each time-point, dependent on a different pattern of neural responses. This would result in a dynamic coding pattern, where object information can be only decoded from classifiers trained and tested at the same time-point, and not when tested at a different time point, because the neural patterns have changed. This would result in a diagonal pattern in the temporal generalisation decoding matrix. In contrast, neural codes could be stable over time, meaning that a classifier trained at one time point can also decode information at other time points, suggesting a shared neural code that generalises over time, indicated by decoding that extends to off-diagonal parts of the matrix. In this way, the temporal generalisation approach can reveal different modes of brain dynamics.

Classifiers were trained using MEG sensor patterns at each time-point after the objects appeared on the screen and tested on new trials. For object shape, despite strong decoding along the diagonal in the first 300 ms, we did not observe any significant clusters in the temporal generalisation matrix (Figure 3a), although a cluster at approximately 100 ms approached significance (cluster p = 0.066). When testing decoding along the diagonal, there were two significant clusters of shape decoding with peak decoding near 100 ms (cluster p < 0.001) and 200 ms (cluster p = 0.010; Figure 3g). For real-world size, there was a significant cluster of decoding in the 300-550 ms time period (cluster p = 0.0002; Figure 3c). This effect was characterised by a dynamic coding pattern, where decoding was significantly greater along the diagonal than the corresponding off-diagonal elements (Oh et al., 2019), followed by a mixed neural code including a stable period of generalisation – when the off-diagonal decoding was both significant and not significantly smaller than the diagonal (Oh et al., 2019) (Figure 3d). For object category, significant decoding was seen in a similar time window, extending to around 600 ms (cluster p = 0.0002; Figure 3e). In contrast to real-world size, the initial dynamic coding pattern was followed by a prominent stable period of generalisation (Figure 3f). This indicates that image-based information about objects is present after approximately 100 ms, while higher-level information about objects, such as real-world size and contextual associations, are present beyond around 200-300 ms.

**Figure 3.**
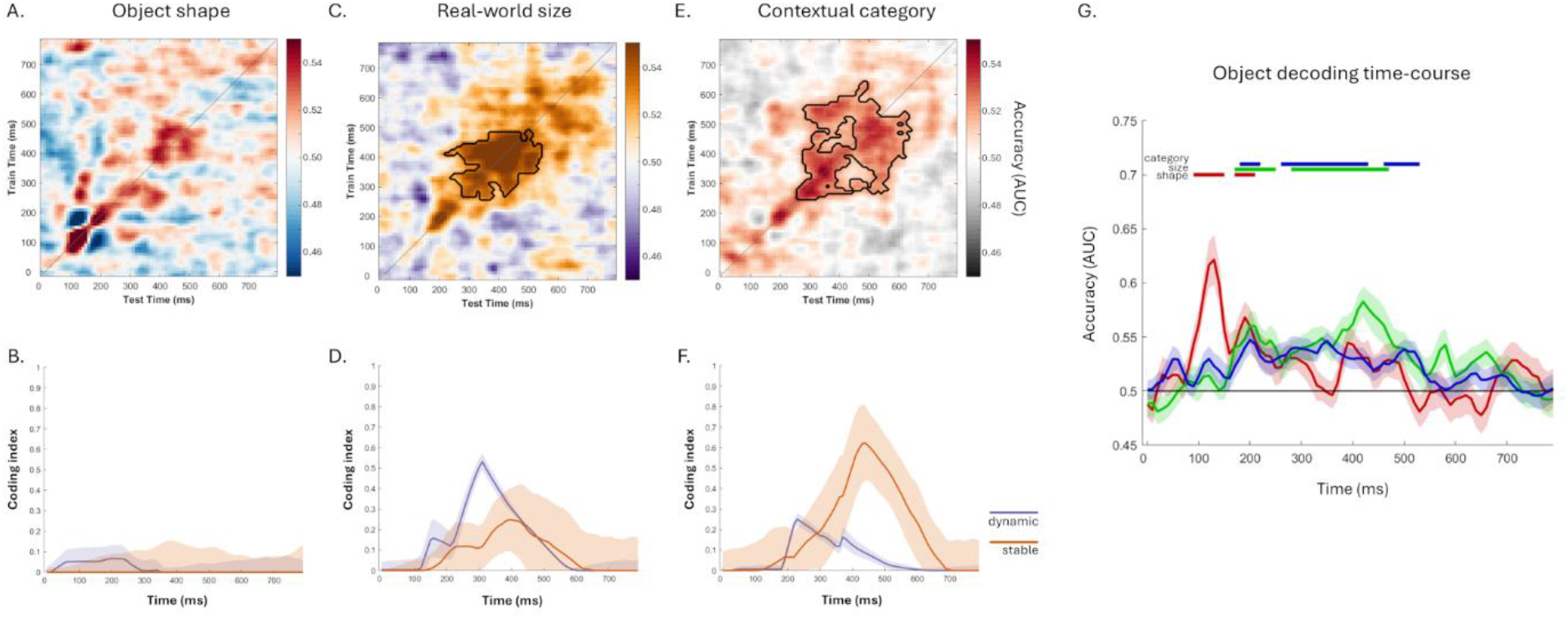
MEG decoding of object information when no context is present. a) Temporal generalisation decoding of object shape. b) Magnitude of dynamic and stable coding patterns over time for classifiers trained to dissociate objects of different shapes. c) Temporal generalisation decoding of real-world size. d) Magnitude of dynamic and stable coding patterns over time for classifiers trained to dissociate real-world size. e) Temporal generalisation decoding of contextual category. f) Magnitude of dynamic and stable coding patterns over time for classifiers trained to dissociate object category. g) Temporally-aligned decoding for each object property. Chance decoding is 0.5. Periods of significant decoding are shown with black outlines in a, c, e and with horizontal lines in g. Shaded areas in b and d show bootstrap 95% CIs, and standard error in g.

### Object information is shifted by context

The previous analyses established that the different forms of information could be decoded from objects without a prior context. Next, we tested whether the timing of object information was changed due to the scene. In other words, is object information shifted in time according to the relationship between the scene and object? To directly address this, we evaluated if object information relating to object shape, real-world size or category is shifted in time compared to a no-context condition. To test this, we first trained classifiers using no context object trials to distinguish between (a) horizontal and vertically orientated objects, (b) big and small objects, and (c) different categories of object. These classifiers were then tested using trials from either congruent or incongruent conditions, to establish the degree, and timing, of any across-condition decoding, and how generalisation differs between congruent and incongruent trials (Figure 2b).

To test for shifts in object information in time, we used a temporal generalisation approach for the congruent and incongruent trials separately. If there was a strong temporal alignment between object information in both the no context trials and scene trials, then we would see significant decoding along the diagonal. If there is a temporal shift in object information, then we would see significant off-diagonal decoding, indicating object information captured from the no-context objects is pushed forward or backwards in time due to the prior scene context. In addition to quantifying stable and dynamic coding patterns, we also tested for shifted decoding patterns by evaluating asymmetries in decoding performance above and below the diagonal axis.

For object shape, classifiers were trained on MEG signals from the no-context objects and tested on the congruent trials. This revealed two significant clusters of decoding which stayed close to the diagonal (Figure 4a), one around 100 to 400 ms (cluster p < 0.001), and a second around 400-500 ms (cluster p = 0.042). Across-condition decoding was predominantly characterised by a dynamic coding pattern (Figure 4b). While both clusters were positioned along the diagonal, we next evaluated if the periods of significant decoding were biased towards the upper or lower halves of the decoding matrix. To test this, we quantified the distance from the diagonal where significant effects were present, creating a distribution. If no bias is present, then this should result in a normal distribution, whereas a bias would present as a deviation from normality. Applying a Jarque-Bera test for normality showed no significant deviation from normality (p = 0.059; Figure 4c). Further evaluation of whether any asymmetry was restricted to specific time periods, we calculated the degree of asymmetry at each time point along the diagonal, again suggesting a lack of bias (Figure 4d). Together this indicates a similar temporal trajectory of object shape information in the no-context and congruent conditions.

**Figure 4.**
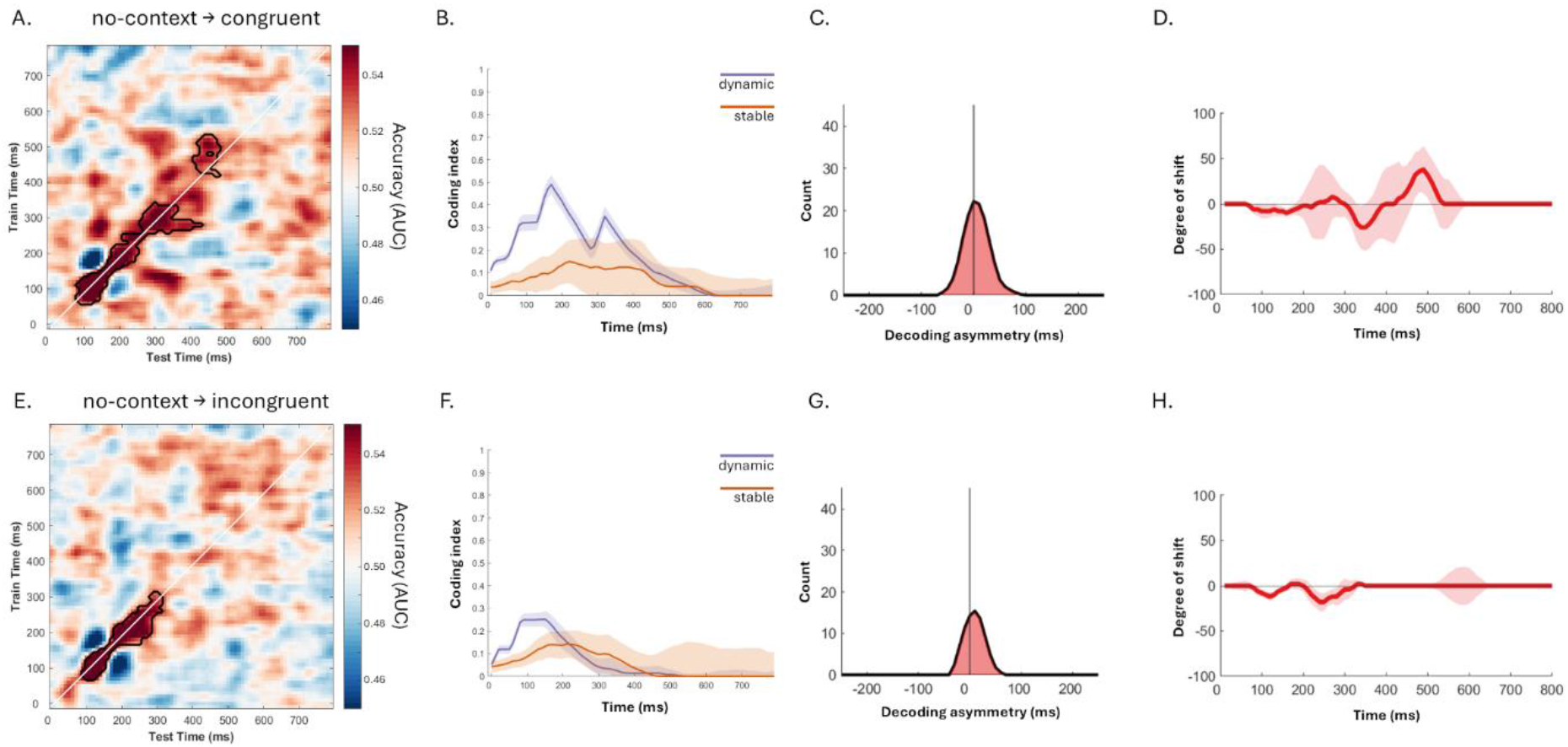
MEG decoding of object shape information across conditions. a) Temporal generalisation decoding accuracy when classifiers are trained on the no-context trials and tested on the congruent trials. b,f) Magnitude of dynamic and stable coding patterns over time. c,g) Distribution of when significant effects occurred relative to the diagonal in the temporal generalisation matrix. d,h) Quantifying the degree of temporal shift in decoding over time. e) Temporal generalisation decoding accuracy when classifiers are trained on the no-context trials and tested on the incongruent trials. In all cases chance decoding is 0.5. Periods of significant decoding are shown with black outlines. Shaded areas show 95% bootstrap CIs.

Training on the no-context objects and testing on the incongruent trials revealed a similar pattern (Figure 4e). A significant cluster was detected along the diagonal around 100-400 ms (cluster p = 0.006; Figure 4e), which was again predominantly linked to a dynamic coding pattern suggesting little generalisation to earlier or later time points (Figure 4f). This was reinforced by the lack of asymmetries in decoding above or below the axis (p = 0.129; Figure 4g), while at no time point did the asymmetry bias substantially deviate from zero (Figure 4h). Together, we observed no evidence of shifted object representations for an image-based measure of shape. However, this would not be expected, as the shape of the image is not predictable given a visual scene. Rather, more semantic properties should be suspectable to contextual influences.

For real-world size, classifiers were trained on MEG signals from the no-context objects and tested on the congruent trials. This revealed two significant clusters of decoding which appeared temporally-aligned (Figure 5a), one around 200 ms (cluster p = 0.043), and a second around 300-600 ms (cluster p = 0.006). Across-condition decoding was characterised by an initial dynamic coding pattern followed by a stable coding pattern (Figure 5b). While both clusters included the diagonal, it also appeared that periods of significant decoding were biased towards the upper half of the decoding matrix. Applying a Jarque-Bera test for normality, we found that the distribution of decoding was significantly different to a normal distribution (p = 0.0109), with more significant periods of generalisations above than below the diagonal (Figure 5c). Calculating this bias at each time point along the diagonal, suggested this shift was most prominent around 400 ms (Figure 5d). Inspection of the temporal generalisation matrix suggested that real-world size information was shifted up to 200 ms earlier following a congruent context (e.g. decoding is significant when training at 500 ms and testing at 300 ms).

**Figure 5.**
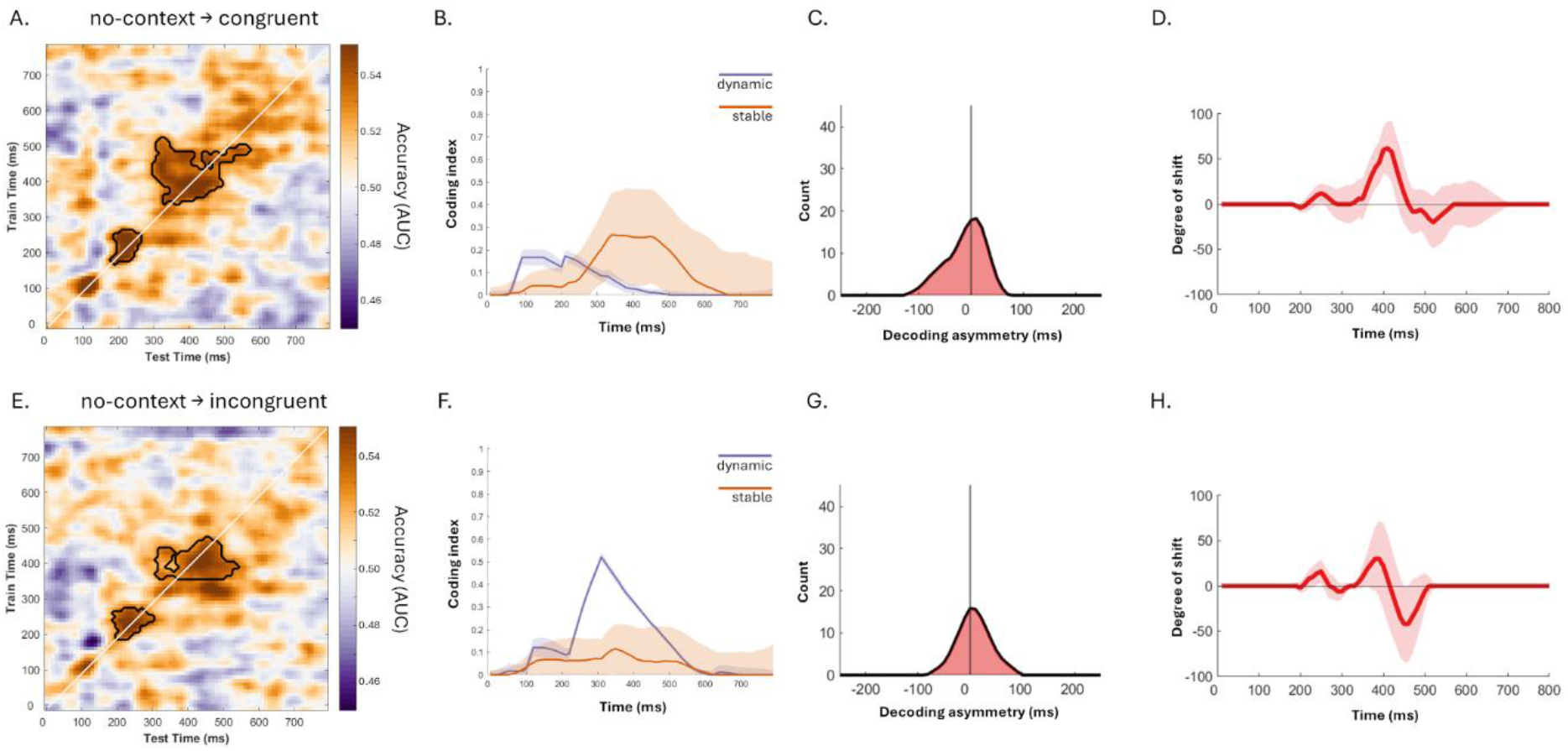
MEG decoding of real-world size information across conditions. a) Temporal generalisation decoding accuracy when classifiers are trained on the no-context trials and tested on the congruent trials. b,f) Magnitude of dynamic and stable coding patterns over time. c,g) Distribution of when significant effects occurred relative to the diagonal in the temporal generalisation matrix, with *c* showing a skewed distribution with more periods of significant decoding shifted earlier in time. d,h) Quantifying the degree of temporal shift in decoding over time. e) Temporal generalisation decoding accuracy when classifiers are trained on the no-context trials and tested on the incongruent trials. In all cases chance decoding is 0.5. Periods of significant decoding are shown with black outlines. Shaded areas show 95% bootstrap CIs.

Training on the no-context objects and testing on the incongruent trials revealed a different pattern. Two significant clusters were detected along the diagonal (Figure 5e), one around 200 ms (cluster p = 0.029), and a second around 300-500 ms (cluster p = 0.0012). For incongruent objects, across-condition decoding was dominated by a dynamic coding pattern suggesting little generalisation to earlier or later time points (Figure 5f). This was reinforced by the lack of asymmetries in decoding above or below the axis (p > 0.5; Figure 5g), while at no time point did the asymmetry bias substantially deviate from zero (Figure 5h). Finally, to assess the possibility of overall enhanced object information in the congruent condition, a direct contrast between congruent and incongruent decoding along the diagonal showed no significant difference in decoding accuracy.

For object category information, training classifiers on MEG signals from the no-context objects and testing on the congruent trials revealed two prominent clusters of significant decoding (Figure 6a). First, a cluster along the diagonal from 100 to 500 ms showing temporally-aligned object representations between the no-context and congruent trials (cluster p < 0.0001). Crucially, a second off-diagonal cluster showing that object information expressed between 600 and 700 ms for the no-context trials was seen earlier in time for the congruent trials, between 400 and 500 ms after the object appeared (cluster p = 0.006). The early temporal alignment followed by off-diagonal decoding is further reflected in the initial prominence of a dynamic coding pattern, followed by an increase in stable coding patterns (Figure 6b). While there is a clear temporal shift in decoding for the off-diagonal cluster, it is also noticeable that decoding overall appears biased towards the upper half of the decoding matrix. Through calculating the degree of bias, it becomes clear that decoding was significantly shifted above the axis (p = 0.0101), with more significant periods of generalisation above than below the diagonal (Figure 6c). Calculating this bias at each time point along the diagonal suggested this shift began around 300 ms to near 700 ms (Figure 6d). Inspection of the temporal generalisation matrix suggested that category information was shifted up to 200 ms earlier following a congruent context.

**Figure 6.**
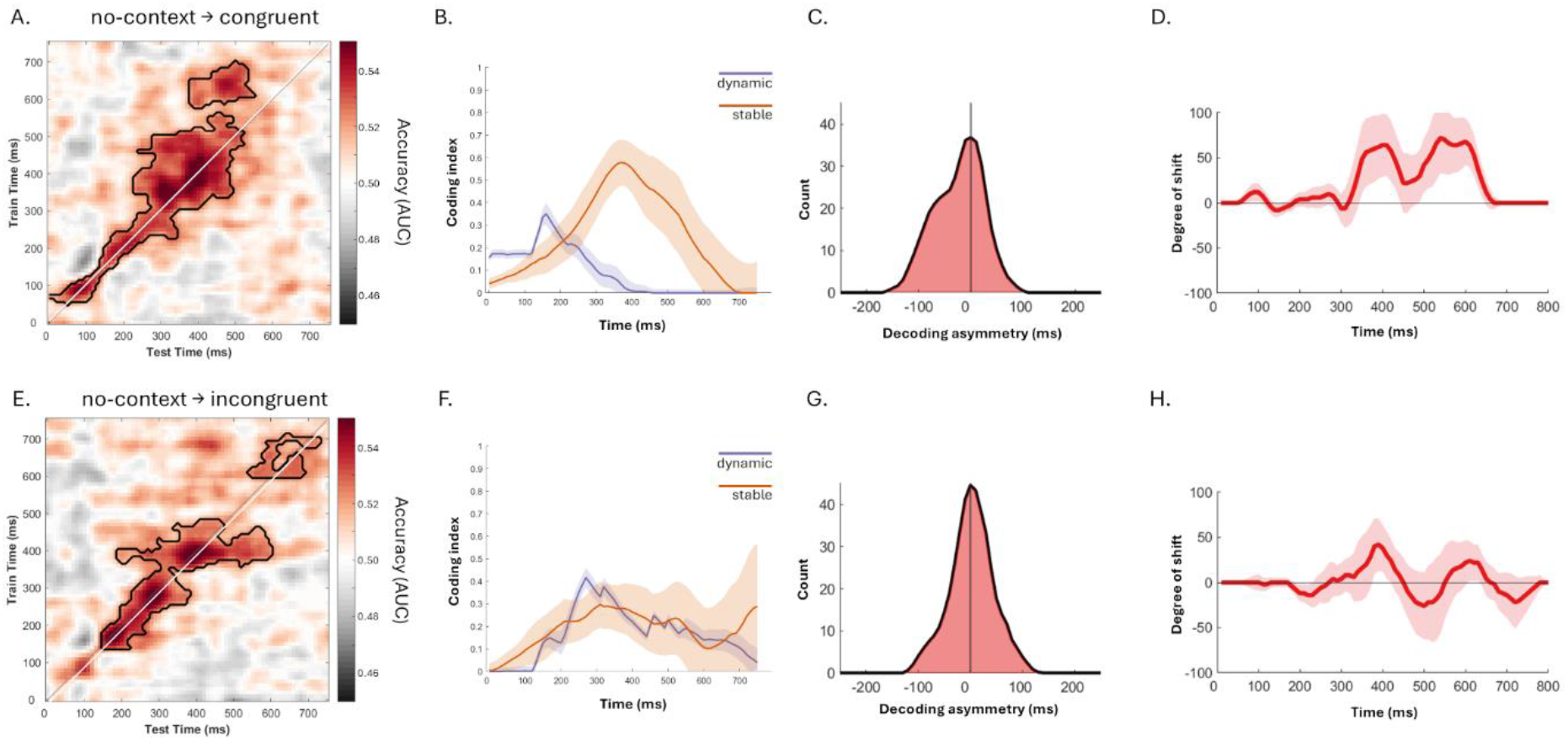
MEG decoding of category information across conditions. a) Temporal generalisation decoding accuracy when classifiers are trained on the no-context trials and tested on the congruent trials. b,f) Magnitude of dynamic and stable coding patterns over time. c,g) Distribution of when significant effects occurred relative to the diagonal in the temporal generalisation matrix, with *c* showing a skewed distribution with more periods of significant decoding shifted earlier in time. d,h) Quantifying the degree of temporal shift in decoding over time. e) Temporal generalisation decoding accuracy when classifiers are trained on the no-context trials and tested on the incongruent trials. In all cases chance decoding is 0.5. Periods of significant decoding are shown with black outlines. Shaded areas show 95% bootstrap CIs.

Training on the no-context objects and testing on the incongruent trials revealed a different pattern (Figure 6e). While two clusters of significant decoding were observed, now both remained close to the diagonal (cluster p < 0.001, p = 0.015), suggesting temporally-aligned category representations for objects in the no-context and incongruent conditions. This was further reflected in the greater proportion of dynamic coding with a less prominent stable coding pattern compared to congruent trials (Figure 6f). Like for real-world size, there was no evidence of asymmetries in decoding above or below the axis (p > 0.5; Figure 6g), while at no point did the bias substantially deviate from zero (Figure 6h). Finally, to assess the possibility of enhanced object information, a direct contrast between congruent and incongruent trials along the diagonal showed no significant difference in decoding accuracy.

Together, our results across both real-world size and contextual category decoding show that object representations occur up to 200 ms earlier in time when the prior scene is congruent with the subsequent object, with effects being restricted to semantic, and not image-based object properties.

### Shifted object information correlates with behavioural facilitation

While the above analyses identified multiple windows of shifted decoding for congruent trials, a salient shift was seen for contextual category decoding where a late cluster, around 600-700 ms, appeared shifted earlier for congruent trials but temporally-aligned for incongruent trials. Closer inspection of decoding in these two clusters using a repeated-measure ANOVA with factors congruency and cluster showed a significant interaction (F=21.82, df = 1,23, p < 0.001; Figure 7a). Follow-up tests showed that the shifted cluster had higher decoding for congruent than incongruent trials (congruent = 53%, incongruent = 51%; t(23) = 1.90, p = 0.035, one-tailed), and the late aligned cluster had higher decoding for incongruent than congruent trials (congruent = 51%, incongruent = 52%; t(23) = 2.26, p = 0.017, one-tailed). To assess if the heightened decoding for congruent trials in the shifted cluster was behaviourally relevant, we correlated the magnitude of the congruency decoding effect with the reaction time congruency effect (Figure 7b). Spearman’s correlation showed a significant negative effect (rho = -0.46, p = 0.026), where better decoding for congruent trials was associated with faster object recognition response times for congruent trials.

**Figure 7.**
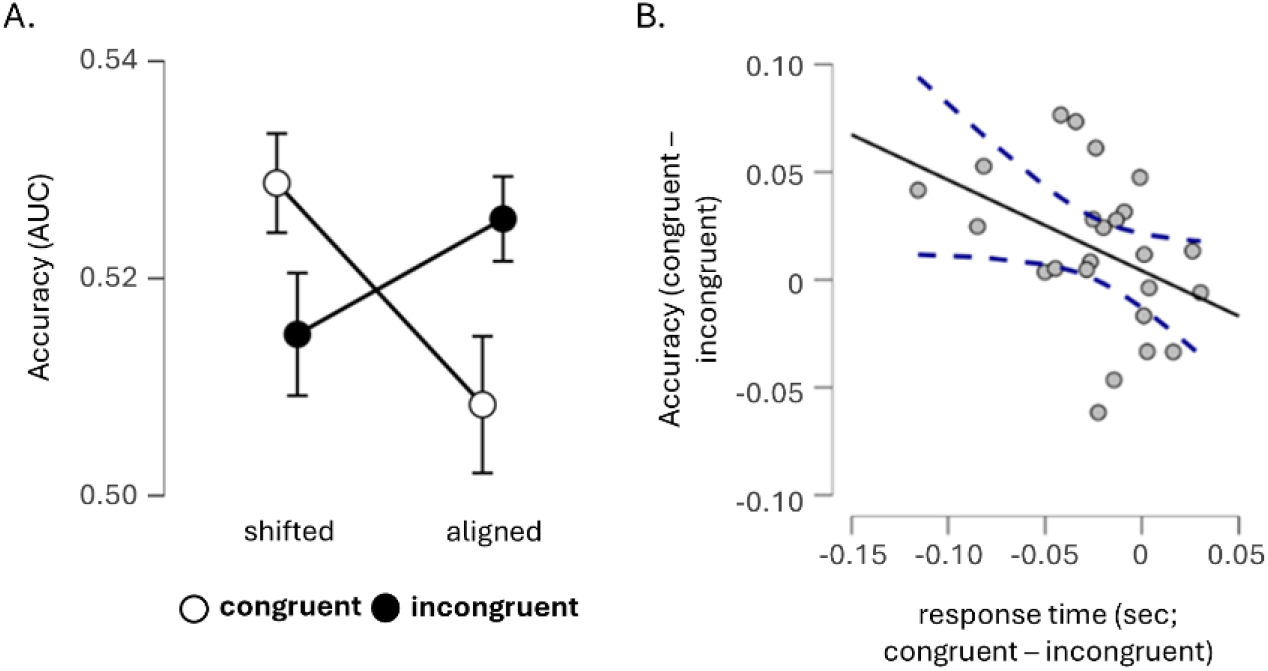
a) Interaction between decoding for the congruent and incongruent trials in the temporally-shifted (off-diagonal) and temporally-aligned (diagonal) clusters. Error bars show standard error of the mean. b) Relationship between the behavioural congruency effect and decoding congruency effect for the shifted (off-diagonal) cluster. Each datapoint is a single participant, with a linear trendline fitted to the data. Dotted lines show 95% confidence intervals.

While this result suggests that the visual scene facilitates object processing such that category information is present earlier in time and correlates with behavioural facilitation, it could also be that decoding differences relate to a maintained representation of the visual scene itself. However, we find this unlikely for two reasons. First, the classifier is trained with data that only knows about objects which were not linked to a prior scene. As such, successful decoding is driven by learned neural patterns following the perception of acontextual objects. Second, if the off-diagonal decoding were driven by a maintained congruent visual scene representation, then decoding would also generalise to time periods when the visual scene was actually present. To test this, classifiers trained using the no-context objects were tested using congruent trials across time periods between the onset of the visual scene and after the object appeared. Decoding during the scene epoch did not show any significant category decoding, which was only seen during the epoch when the object was present (Figure 8). This confirms that the classifier training picked up on categorical object representations which do not generalise to viewing different scene types.

**Figure 8.**
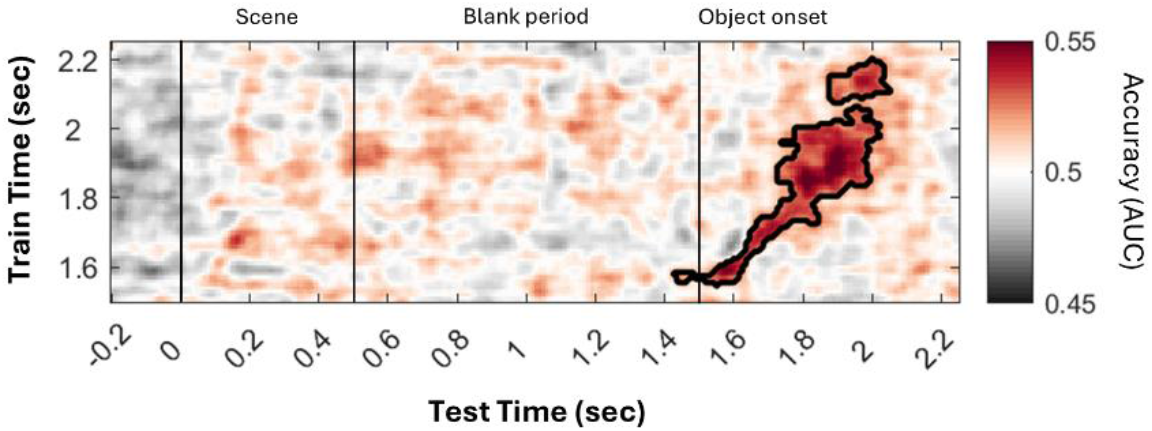
Temporal generalisation for the congruent trials across the scene, delay and object periods.

Together, our results suggest that semantic object processing is shifted forward in time when prior scene information provides appropriate expectations for the kind of object that might occur next, and that this temporal shift in categorical object representations facilitates the recognition of objects.

## Discussion

The present research clearly demonstrates that our visual environment can dynamically warp the temporal trajectory of object recognition, and accelerate the progression through representational states that support recognition. Using MEG and temporal generalisation decoding, we showed that object information is shifted up to 200 ms earlier when the preceding visual scene was congruent with the object. This effect was observed for two distinct forms of semantic object information – an objects real-world size and its contextual category, while the strength of the category effect was associated with behavioural facilitation. These findings support the hypothesis that the prior visual scene results in a temporal shift in object information, where mnemonic information related to the scene might act to accelerate object information, making recognition faster and more efficient. Moreover, this suggests that visual object recognition is not an immutable bottom-up sensory driven process, but is a flexible inferential process conditioned on our current context.

A large body of behavioural and neuroimaging research shows that an objects’ context influences how objects are processed and the decisions people make. Using time-sensitive brain imaging techniques, numerous studies show that context modulates processing between 200 and 600 ms after seeing an object, with most interpretations relating to accessing semantic knowledge, integration of the object and context, or an expectation violation (Brandman & Peelen, 2017; Coco et al., 2020; Ganis & Kutas, 2003; Gu et al., 2025; Kumar et al., 2021; Kutas & Federmeier, 2011; Lauer et al., 2018; Mudrik et al., 2014; Võ & Wolfe, 2013). The prevailing interpretation is that the strength of these processes is changed, but with an otherwise fixed temporal sequence. Our findings reveal a new way to conceptualise how context impacts object recognition, where context can dynamically warp the temporal trajectory of object processing. This means that previous research highlighting modulations between conditions, might actually reflect representational shifts in time and a temporal divergence between conditions, that is only observed by the representational time-course of each condition.

The central finding in this research is that neural patterns related to real-world size and contextual category were shifted earlier in time by a preceding congruent scene. This was revealed using across-condition temporal generalisation decoding, where classifiers were trained on objects without a context, and tested on the same objects seen after a congruent or incongruent visual scene. We observed clear shifts in decoding above the diagonal axis indicating that object information was present earlier in time when objects were preceded by a congruent scene. This pattern was not observed for image-based object properties, which can not be predicted from the scene, or for the incongruent scenes, which remained temporally aligned with no-context objects. Further, we did not find strong evidence that congruent scenes simply enhanced the decodability of objects but without temporal a shift.

The temporal generalisation decoding results add new dimensions to previous decoding research. Using EEG, Chen et al., (2022) found that object decoding was enhanced for congruent compared to incongruent trials. While we do see some evidence in line with this, our primary effects were shown through a bias above the decoding axis, suggesting that context might have a wider role, where both enhancement and a temporal warping can occur.

Our evidence of earlier object information for congruent compared to incongruent trials expands on past EEG evidence. Truman and Mudrik (2018) presented congruent and incongruent scene-object images, in addition to scene images where the object was replaced with a scrambled item, making object recognition impossible. They reported that EEG signals for congruent trials diverged from scrambled trials earlier than incongruent trials diverged from the scrambled trials. This was seen as evidence of more rapid object identification when objects are congruent with the scene they appeared within. Further, using neural pattern similarity analysis of EEG data, we previously showed that semantic information associated with objects was resolved earlier in time when objects were seen within a congruent scene compared to an incongruent scene (Krugliak et al., 2023). The current research builds on these findings by showing that context can shift the temporal trajectory of object representations, impacts multiple dimensions of object knowledge, and facilitates behaviour.

Our novel results add important new insights to accounts of how object and scene processing interact. It is well established that a visual scene can modulate the processing of objects, where a contextual facilitation occurs (Oliva & Torralba, 2007; Peelen et al., 2024; Trapp & Bar, 2015). While such facilitation might be related to predictive interactions between object and scene networks (Peelen et al., 2024), investigations into whether the timing of object processing is shifted through contextual facilitation are hampered by paradigms that don’t capture the unfolding nature of real events. In real-life, our environmental context is relatively stable over time while objects are relatively transient. Walking down the street, we may have global expectations about the kinds of things we see, and different items will come and go from view. As such, contextually-mediated expectations are about the future, and will modulate the recognition process of objects after they come into view.

Such temporal uncoupling between expectations and the following perceptual experience has been demonstrated through fMRI, TMS, and MEG. Using paradigms where a cue is predictive of a later stimulus, past research indicates that hippocampal patterns pre-activate the expected perceptual details in visual cortex when a specific visual image can be predicted (Clarke et al., 2022; Hindy et al., 2016). Using MEG, it is suggested that expectations can enhance the representation of predictable motion patterns after around 150 ms (Aitken et al., 2020), while applying TMS to category-specific regions during a cue period extinguishes expectations-related behavioural benefits when seeing subsequent images (Gandolfo & Downing, 2019). Paradigms using simultaneous object-scene presentations could therefore minimise this vital predictive dynamic, which we incorporate by using a preceding scene image, followed by an object. This allows scene-generated predictions to act on future objects. Within this predictive framework, we argue that expectations about schema-consistent events can act to accelerate the recognition process and facilitate behaviour.

This raises a further important question. What do expectations *do* to neural populations? It has been suggested that top-down information can sharpen object representations, such that feedback enhances the representation of expected objects and supresses unexpected objects (de Lange et al., 2018; Peelen et al., 2024; Trapp & Bar, 2015; Yon et al., 2018). In our paradigm, any expectations generated by the scene will be at a semantic level and relate to a large array of schema-consistent items, and not visual details of images. As such, expectation effects would not be expected at the representational level of visual stimulus features, and to occur within the typical time frame when semantic object properties are accessed. This is indeed what we observed, with a representational shift occurring for multiple stimulus features requiring access to high-level semantics (real-world size and contextual category) and occurring beyond 300 ms, which were not present for image-based stimulus features. This is consistent with accounts where scene-derived expectations constrain the range of likely object identities (Bar, 2004; Trapp & Bar, 2015) and thereby facilitate access to semantic object representations. While we did not identify the neural generators of our effects, our past work on object semantics and context suggest the involvement of the posterior ventral temporal cortex and anterior temporal lobes (Gu et al., 2025; von Seth et al., 2023).

Finally, if the neural mechanisms of object recognition are conditional on the preceding context, and we would argue all real-world objects are perceived in the context of the recent past, present and anticipated future, then traditional models of object recognition might be limited for real-world applications. More broadly, our findings suggest that object processing occurs as a trajectory through representational states, which our prior knowledge can flexibly alter resulting in representations emerging at different times.

## Methods

### Participants

Twenty-four participants (15 females, age range 18-35 years) took part in the MEG study. All were right-handed and had normal or corrected-to-normal vision, and gave written informed consent. The study was approved by the Cambridge University Psychology Research Ethics Committee (PRE.2019.051).

### Stimuli

A total of 450 colour images were used in the study, including 150 scene images, 150 scrambled scene images, and 150 object images. The scenes images were obtained from Lauer et al. (2018) the SUN397 scene image database (Xiao et al., 2010), and internet searches using Google Image Search. The scene images were of a household location (n=50), an urban street view (n=50) or of a rural location (n=50). Objects were shown in colour, isolated on a white background, from the Hemera photo object image set or Clarke et al. (2018).

Each scene image was paired with two object images, one that was congruent with the scene, and one that was incongruent (Figure 1a). Objects only appeared once with a congruent scene and once with an incongruent scene, meaning that each scene was paired with two objects, and each object was paired with two scenes. This resulted in 150 congruent pairs and 150 incongruent pairs where each scene and object appeared once in each list. Scrambled scene images were created from the two scene images paired with that object. The two scene images were divided into small squares and a new image was created by combining half the squares from one scene with half from the other scene. The squares were flipped and randomly shuffled to create a scrambled image, resulting in 150 scrambled (no content) images.

### Paradigm

Each trial displayed a scene image (or scrambled image) followed by an object. Each trial began with a fixation cross lasting 750 ms before a scene (or scrambled) image was shown for 500 ms. This was followed by a blank grey screen lasting 1 second. An object was then displayed in the centre of the screen for 50 ms followed by a dynamic backwards mask lasting 200 ms (Figure 1b).

Participants pressed a button using their right index finger if they recognized the object and knew what the object was, or using their right middle finger if they did not recognize the object. For objects that were recognized, they next answered a yes/no question about the object’s characteristics (e.g., ‘‘Is the object edible?’’, ‘‘Can you buy the object in a supermarket?’’), and was used to facilitate a deeper processing of the object and to discourage participants to rely on visual features alone. This was followed by a congruency rating for the object and scene pair (this was omitted if the scene was a scrambled image). Congruency ratings were made using the left hand to the question ‘How likely would you expect to encounter this object in this context?’, with 3 response options (unlikely, neutral, likely).

Each object was shown three times, on each occasion being preceded by either the congruent, incongruent or scrambled scene image, with a total of 450 trials, which were presented in a randomized order. The experiment was presented on a screen positioned 120 cm from the participant with a resolution of 1024×768. All scene images were full screen, and object images had a resolution of 500×500 pixels. The experiment was controlled using MATLAB (MathWorks, Natick, MA) and Psychtoolbox (Brainard, 1997).

### Behavioural analysis

For each participant, we collected object recognition reaction times and the rated congruency between the object and the preceding scene. Reaction times for successful recognition trials were classified into congruent or incongruent based on the participants own ratings, or the no context condition if the object was preceded by a scrambled scene. Repeated measures ANOVA and follow-up tests were run using JASP 0.18.3.0., and follow-up t-tests were corrected for multiple comparisons using the Holm-Bonferroni correction.

### MEG acquisition

Continuous MEG data were recorded using a whole-head 306 channel (102 magnetometers, 204 planar gradiometers) TRIUX neo system (MEGIN, Espoo, Finland) at the Cognition and Brain Sciences Unit, Cambridge, UK. Five head-position indicator (HPI) coils were used to record the head position every 200 ms within the MEG helmet, and eye movements and blinks were monitored using the electrooculogram (EOG) recorded bipolarly through electrodes placed above and below the right eye (vertical) and at the outer canthi (horizontal). The participants’ head shape was digitally recorded using a 3D digitizer (Fastrak Polhemus, Inc., Colchester, VA, USA) along with the positions of the EOG electrodes, HPI coils, and fiducial points (nasion, left and right periarticular). MEG signals were recorded at a sampling rate of 1000 Hz, with a high-pass filter of 0.03 Hz.

### MEG processing

The MEG data were processed using MNE-python (https://mne.tools/). The raw signals were first Maxfiltered, where static bad channels were detected and subsequently reconstructed by interpolating neighbouring channels, as were bad channels containing long periods of high amplitude or noisy signals. The temporal extension of the signal-space separation (SSS) technique was applied to the data every 4s to segregate the signals originating from within the participants’ heads from those generated by external sources of noise, along with head movement compensation and transformation of the head position to a common position.

The data were then down-sampled to 200 Hz. For Independent Component Analysis (ICA), the data were first high-pass filtered at 1 Hz, and then epoched around the onset of the scenes from -200 ms to 2000 ms, which also includes the object presentation. Epochs where the amplitude exceeded 5e-12 for magnetometers and 4000e for gradiometers were automatically rejected. ICA was computed using ‘fastica’ method, with 60 components, and fitted to the epoched data. ICA components representing EOG and ECG components were detected automatically with MNE built-in functions (ica.find_bads_eog and ica.find_bads_ecg). Where automatic detection failed, the components were identified and excluded manually. (we also rejected some components that represented magnetic artifacts, based on visual inspection). For the main analysis, the downsampled data were filtered between 0.3 Hz and 40 Hz, and epoched around the onset of the object from -200 ms to 800 ms. Noisy epochs were identified and rejected based on visual inspection. No baseline correction was applied (because the blank period between the scene and object allowed for expectations about what objects might appear next to build up and be related to later object processing, and is as such not a traditional baseline period). Finally, ICA was applied to the epoched data.

### Decoding analyses: No-context condition

We performed decoding related to three different object dimensions – object shape, real-world size, and the contextual category of the object.

Object shape decoding distinguished between images where the main object axis was horizontal (n = 52) and those where the main axis was vertical (n = 49). Objects with equal horizontal and vertical proportions were excluded from decoding (e.g. an apple; n = 49). To establish whether MEG signals distinguished between horizontal and vertically orientated images within the no-context condition, we performed a series of temporal generalisation decoding analyses using MNE and sklearn. The preprocessed MEG signals from all 302 channels were used. Logistic regression classifiers were used to discriminate between big and small objects using 10-fold cross-validation. The data used in the training sets was set to be balanced. Classifiers were trained at each time point between the onset of the object image and 800 ms post-object, and evaluated at all time points. Classification accuracy was then averaged across all folds.

Real-world size decoding distinguished between big and small objects based on our knowledge of those items, and not based on the image size. Small objects were defined as those smaller than a brick (n = 56), and large objects as bigger than a bathroom sink (n = 55). Objects not falling into these groups were excluded from decoding (n = 39). Logistic regression classifiers were used to discriminate between big and small objects using 10-fold cross-validation, and averaged across all folds.

For contextual category decoding, trials were divided based on whether the objects were household objects (n = 50), street objects (n = 50) or rural objects (n = 50). Logistic regression classifiers were used to discriminate between each pair of categories (e.g. urban vs rural objects) using 10-fold cross-validation, and averaged across all folds and across the three pairs of categories.

### Decoding analyses: Across conditions

Critically, we evaluated the performance of classifiers trained on the no-context condition against conditions when the object was preceded by a scene (i.e. congruent trials or incongruent trials) to determine if representations were shared across conditions, and whether the decoding was temporally-aligned – meaning decoding was present when training and testing time were the same, or temporally-shifted – meaning decoding was present at time-point preceding or following the time point at which the classifier was trained.

Logistic regression classifiers were trained to discriminate between a pair of categories (e.g. horizontal vs vertical; big vs small; urban vs rural objects etc) for the no-context trials, and tested using trials of the same type from the congruent and incongruent conditions separately. Classifiers were trained at each time point between the onset of the object image and 800 ms post-object, and tested at all time points.

### Cluster-based statistics

Statistically significant periods of decoding were assessed using cluster-based permutation testing (Maris & Oostenveld, 2007). One-sample t-tests using an alpha of 0.01 were used at each element of the temporal generalisation matrix to assess if decoding was above chance (0.5), and t-values were summed within the observed contiguous clusters. Permutation testing was used to evaluate the significance of these clusters. Data for each participant was randomly flipped around the chance axis before one-sample t-tests of the permuted data with the largest cluster size being retained. This was repeated 10,000 times, creating a null distribution of cluster sizes. The cluster p value for the observed clusters was defined as the proportion of permutation-based cluster sizes (which included the observed cluster sizes) that was greater than the observed clusters.

### Stable and dynamic coding

We additionally quantified the proportions of stable and dynamic coding patterns seen in the temporal generalisation matrix (Oh et al., 2019). This analysis seeks to visualise the proportion of significant decoding that primarily occurs along the diagonal that does not generalise over time (dynamic coding), compared to decoding that generalises over time away from the diagonal (stable decoding).

Dynamic coding at each time point was calculated by first testing whether decoding was significantly greater (p < 0.01 uncorrected) along the diagonal compared to generalisation points parallel to the diagonal. We then assess the proportion of elements that showed higher decoding along the diagonal compared to the generalisation area within an area centred at each time-point. Specifically, we created a square on the diagonal at time-point *t* with 200ms width/height and quantified the proportion of those cells where the diagonal decoding was greater than the off-diagonal elements. To avoid the influence of data close to time *t*, data elements were excluded within 30 ms of *t*. Conversely, stable decoding was quantified by first establishing when off-diagonal decoding was not significantly less than diagonal decoding, while both diagonal and off-diagonal decoding were themselves significantly better than chance. Like above, the proportion of stable decoding was also assessed within a moving window over time, with windows centred on the diagonal. Bootstrap 95% confidence intervals were obtained for each coding index, using 1000 random samples of participant data with replacement.

### Asymmetries in decoding

The temporal generalisation approach not only allows for determining the relative dynamic or stable coding pattern, but can also reveal if neural patterns that drive training share representations at a different point in time in the testing data (King & Dehaene, 2014). Therefore decoding above the diagonal indicates a bias towards an earlier time in the test data than training data, and a decoding shifted below the axis indicates a bias later in time in the test data compared to the training data (Figure 2b).

In our across-condition decoding, we sought to quantify this bias to test if object information when no context is present is shifted in time when a prior context is introduced. A global measure of asymmetry was calculated by counting the number of significant elements in the temporal generalisation matrix at the diagonal and at each of the off-diagonal axes. This results in a distribution showing how many significant elements occurred at each distance from the diagonal. To test if this distribution is biased and not symmetric around the diagonal, a Jarque-Bera test for normality was applied. We also calculated a time-sensitive measure of asymmetry, by counting the number of significant effects perpendicular to each time-point along the diagonal. Comparing the number of significant effects above and below the diagonal therefore gives us a time-sensitive estimate of any bias. Variance of this bias was estimated with 95% bootstrap confidence intervals using 1000 random samples of participant data with replacement.

## Acknowledgments

This research was funded in whole, or in part, by the Wellcome Trust (grant no. 211200/Z/18/Z to AC) and the Isaac Newton Trust. For the purpose of open access, the author has applied a CC BY public copyright licence to any Author Accepted Manuscript version arising from this submission.

The authors declare no conflicts of interest.

## References

Aitken, F., Turner, G., & Kok, P. (2020). Prior Expectations of Motion Direction Modulate Early Sensory Processing. The Journal of Neuroscience: The Official Journal of the Society for Neuroscience, 40(33), 6389–6397. 10.1523/JNEUROSCI.0537-20.2020

Bankson, B. B., Hebart, M. N., Groen, I. I. A., & Baker, C. I. (2018). The temporal evolution of conceptual object representations revealed through models of behavior, semantics and deep neural networks. NeuroImage, 178, 172–182. 10.1016/j.neuroimage.2018.05.037

Bar, M. (2004). Visual objects in context. Nat Rev Neurosci, 5(8), Article 8. (15263892). 10.1038/nrn1476

Biederman, I., Mezzanotte, R. J., & Rabinowitz, J. C. (1982). Scene perception: Detecting and judging objects undergoing relational violations. Cognitive Psychology, 14(2), 143–177. 10.1016/0010-0285(82)90007-x

Brandman, T., & Peelen, M. V. (2017). Interaction between scene and object processing revealed by human fMRI and MEG decoding. Journal of Neuroscience, 0582–17. 10.1523/JNEUROSCI.0582-17.2017

Carlson, T., Tovar, D. A., Alink, A., & Kriegeskorte, N. (2013). Representational dynamics of object vision: The first 1000 ms. Journal of Vision, 13(10), Article 10. 10.1167/13.10.1

Chen, L., Cichy, R. M., & Kaiser, D. (2022). Semantic scene-object consistency modulates N300/400 EEG components, but does not automatically facilitate object representations. Cerebral Cortex, 32(16), 3553–3567. 10.1093/cercor/bhab433

Cichy, R. M., Pantazis, D., & Oliva, A. (2014). Resolving human object recognition in space and time. Nature Neuroscience, 17(3), Article 3. 10.1038/nn.3635

Clarke, A., Crivelli-Decker, J., & Ranganath, C. (2022). Contextual Expectations Shape Cortical Reinstatement of Sensory Representations. Journal of Neuroscience, 42(30), Article 30. 10.1523/JNEUROSCI.2045-21.2022

Clarke, A., Devereux, B. J., Randall, B., & Tyler, L. K. (2015). Predicting the Time Course of Individual Objects with MEG. Cerebral Cortex, 25(10), Article 10. 10.1093/cercor/bhu203

Clarke, A., Devereux, B. J., & Tyler, L. K. (2018). Oscillatory Dynamics of Perceptual to Conceptual Transformations in the Ventral Visual Pathway. Journal of Cognitive Neuroscience, 30(11), Article 11. 10.1162/jocn_a_01325

Clarke, A., Taylor, K. I., Devereux, B., Randall, B., & Tyler, L. K. (2013). From Perception to Conception: How Meaningful Objects Are Processed over Time. Cerebral Cortex, 23(1), Article 1. 10.1093/cercor/bhs002

Coco, M. I., Nuthmann, A., & Dimigen, O. (2020). Fixation-related Brain Potentials during Semantic Integration of Object–Scene Information. Journal of Cognitive Neuroscience, 32(4), 571–589. 10.1162/jocn_a_01504

Davenport, J. L., & Potter, M. C. (2004). Scene consistency in object and background perception. Psychological Science, 15(8), Article 8. 10.1111/j.0956-7976.2004.00719.x

de Lange, F. P., Heilbron, M., & Kok, P. (2018). How Do Expectations Shape Perception? Trends in Cognitive Sciences, 22(9), Article 9. 10.1016/j.tics.2018.06.002

Eddine, S., Brothers, T., Wang, L., Spratling, M., & Kuperberg, G. R. (2024). A predictive coding model of the N400. Cognition, 246, 105755. 10.1016/j.cognition.2024.105755

Gandolfo, M., & Downing, P. E. (2019). Causal Evidence for Expression of Perceptual Expectations in Category-Selective Extrastriate Regions. Current Biology: CB, 29(15), 2496–2500.e3. 10.1016/j.cub.2019.06.024

Ganis, G., & Kutas, M. (2003). An electrophysiological study of scene effects on object identification. Cognitive Brain Research, 16(2), Article 2. 10.1016/S0926-6410(02)00244-6

Gu, Y., Krugliak, A., & Clarke, A. (2025). Neural oscillations and top-down connectivity are modulated by object-scene congruency. Cerebral Cortex, 35(10), bhaf290. 10.1093/cercor/bhaf290

Henderson, J. M., & Hollingworth, A. (1999). High-Level Scene Perception. Annual Review of Psychology, 50, 243–271. 10.1146/annurev.psych.50.1.243

Hindy, N. C., Ng, F. Y., & Turk-Browne, N. B. (2016). Linking pattern completion in the hippocampus to predictive coding in visual cortex. Nature Neuroscience, 19(5), 665–667. 10.1038/nn.4284

Joubert, O. R., Rousselet, G. A., Fize, D., & Fabre-Thorpe, M. (2007). Processing scene context: Fast categorization and object interference. Vision Research, 47(26), Article 26. 10.1016/j.visres.2007.09.013

Jozwik, K. M., Kietzmann, T. C., Cichy, R. M., Kriegeskorte, N., & Mur, M. (2023). Deep Neural Networks and Visuo-Semantic Models Explain Complementary Components of Human Ventral-Stream Representational Dynamics. The Journal of Neuroscience, 43(10), 1731–1741. 10.1523/JNEUROSCI.1424-22.2022

Jozwik, K. M., Kriegeskorte, N., & Mur, M. (2016). Visual features as stepping stones toward semantics: Explaining object similarity in IT and perception with non-negative least squares. Neuropsychologia, Special Issue: Functional Selectivity in Perceptual and Cognitive Systems - A Tribute to Shlomo Bentin (1946-2012), 83, 201–226. 10.1016/j.neuropsychologia.2015.10.023

Kaiser, D., Azzalini, D. C., & Peelen, M. V. (2016). Shape-independent object category responses revealed by MEG and fMRI decoding. Journal of Neurophysiology, 115(4), 2246–2250. 10.1152/jn.01074.2015

King, J.-R., & Dehaene, S. (2014). Characterizing the dynamics of mental representations: The temporal generalization method. Trends in Cognitive Sciences, 18(4), Article 4. 10.1016/j.tics.2014.01.002

Krugliak, A., Draschkow, D., Võ, M. L.-H., & Clarke, A. (2023). Semantic object processing is modulated by prior scene context. Language, Cognition and Neuroscience, 0(0), 1–10. 10.1080/23273798.2023.2279083

Kumar, M., Federmeier, K. D., & Beck, D. M. (2021). The N300: An Index for Predictive Coding of Complex Visual Objects and Scenes. Cerebral Cortex Communications, 2(2), Article 2. 10.1093/texcom/tgab030

Kutas, M., & Federmeier, K. D. (2011). Thirty years and counting: Finding meaning in the N400 component of the event-related brain potential (ERP). Annual Review of Psychology, 62, 621–647.

Lauer, T., Cornelissen, T. H. W., Draschkow, D., Willenbockel, V., & Võ, M. L.-H. (2018). The role of scene summary statistics in object recognition. Scientific Reports, 8(1), Article 1. 10.1038/s41598-018-32991-1

Leonardelli, E., & Fairhall, S. L. (2022). Similarity-based fMRI-MEG fusion reveals hierarchical organisation within the brain’s semantic system. NeuroImage, 259, 119405. 10.1016/j.neuroimage.2022.119405

Maris, E., & Oostenveld, R. (2007). Nonparametric statistical testing of EEG- and MEG data. Journal of Neuroscience Methods, 164, 177–190.

Mudrik, L., Shalgi, S., Lamy, D., & Deouell, L. Y. (2014). Synchronous contextual irregularities affect early scene processing: Replication and extension. Neuropsychologia, 56, 447–458. 10.1016/j.neuropsychologia.2014.02.020

Oh, B.-I., Kim, Y.-J., & Kang, M.-S. (2019). Ensemble representations reveal distinct neural coding of visual working memory. Nature Communications, 10(1), 5665. 10.1038/s41467-019-13592-6

Oliva, A., & Torralba, A. (2007). The role of context in object recognition. Trends in Cognitive Sciences, 11(12), Article 12. 10.1016/j.tics.2007.09.009

Palmer, Stephen E. (1975). The effects of contextual scenes on the identification of objects. Memory & Cognition, 3(5), Article 5. 10.3758/BF03197524

Peelen, M. V., Berlot, E., & de Lange, F. P. (2024). Predictive processing of scenes and objects. Nature Reviews Psychology, 3, 13–26. 10.1038/s44159-023-00254-0

Press, C., Kok, P., & Yon, D. (2020). The Perceptual Prediction Paradox. Trends in Cognitive Sciences, 24(1), 13–24. 10.1016/j.tics.2019.11.003

Trapp, S., & Bar, M. (2015). Prediction, context, and competition in visual recognition. Annals of the New York Academy of Sciences, 1339(1), Article 1. 10.1111/nyas.12680

Truman, A., & Mudrik, L. (2018). Are incongruent objects harder to identify? The functional significance of the N300 component. Neuropsychologia, 117, 222–232. 10.1016/j.neuropsychologia.2018.06.004

Võ, M. L.-H. (2021). The meaning and structure of scenes. Vision Research, 181, 10–20. 10.1016/j.visres.2020.11.003

Võ, M. L.-H., & Wolfe, J. M. (2013). Differential Electrophysiological Signatures of Semantic and Syntactic Scene Processing. Psychological Science, 24(9), Article 9. 10.1177/0956797613476955

von Seth, J., Nicholls, V. I., Tyler, L. K., & Clarke, A. (2023). Recurrent connectivity supports higher-level visual and semantic object representations in the brain. Communications Biology, 6(1), 1–15. 10.1038/s42003-023-05565-9

Xiao, J., Hays, J., Ehinger, K. A., Oliva, A., & Torralba, A. (2010). SUN database: Large-scale scene recognition from abbey to zoo. 2010 IEEE Computer Society Conference on Computer Vision and Pattern Recognition, 3485–3492. 10.1109/CVPR.2010.5539970

Yon, D., Gilbert, S. J., de Lange, F. P., & Press, C. (2018). Action sharpens sensory representations of expected outcomes. Nature Communications, 9(1), 4288. 10.1038/s41467-018-06752-7

